# A New Model of Genetic Variation and Evolution Evaluates Relative Impacts of Background Selections and Selective Sweeps

**DOI:** 10.1101/2020.01.11.901066

**Authors:** Xun Gu

## Abstract

Intra-population genetic variation and interspecies divergence in chromosome regions can be considerably affected by different local recombination rates. There are two models: (*i*) the selective sweeps that reduces the genetic diversity at linked sites and elevates the divergence rate; and (*ii*) the background selection that reduces the genetic diversity at linked sites and divergence rate. An intriguing question, yet highly controversial, is which one is dominant. In this paper, I develop a framework of generalize background selection, formulated by a diffusion model with two killing functions: the one associated with (negative) background selection is the rate to stop a fixation process of a mutation randomly, and the other associated with positive background selection (selective sweep) is the rate to stop a loss process of a mutation randomly. A simple relationship between the level of reduced diversity and the rate of divergence is derived, depending on the strength of generalized background selection (*G*) and the proportion of positive background selection (*β*). We analyzed the interspecies divergence and intra-population diversity in low-recombination regions of three organisms (fruitfly, soybean and human). Strikingly, all datasets demonstrated the dominance of (negative) background selection, and the positive background selection (selective sweeps) only has a small contribution (*β∼*10%). However, our analysis rejects the notion of *β=0*, namely, a complete negative background selection is unlikely. These findings may shed some lights on the long-term debates around Neutral Theory.

## Introduction

Levels of intra-species genetic variation and rates of molecular evolution in different regions of the genome may be greatly affected by differences in the rate of recombination (1, 2). Maynard Smith and Haigh (3) proposed the hitchhiking hypothesis, where the spread of a favorable mutation reduces the level of neutral or nearly-neutral variability at linked sites; a process also termed ‘selective sweeps’ (4). On the other hand, Charlesworth et al. (5) argued that reduction of genetic variation in low-recombination regions can operate through the removal effects of purifying selections on deleterious mutations, a process called ‘background selections’. One may see a number of comprehensive reviews for a rich body of literatures in both theoretical an empirical studies (6-11).

The advent of high throughput genomics and the long-standing selectionist-neutralist debate (12, 13) have inspired tremendous attempts to infer the relationship between the level of intra-species diversity, the local recombination rate and the interspecies divergence (14, 15). Pioneered by Kaplan et al. (16), Wiehe and Stephan (17) and Kim and Stephan (18), most studies concluded that considerable reduction in genetic diversity was the result of joint effects of selective sweeps and background selections (19-23). By contrast, low-recombination chromosome regions generally showed no considerable reduced or elevated interspecies divergence (24-27). Because background selection predicts a much lower interspecies divergence in low-recombination regions whereas selective sweep predicts a much higher one, this observation has been widely interpreted as the existence of selective sweeps under the background selections. Therefore, an intriguing question is which of these two nonexclusive causal factors is more dominant (7, 28, 29); yet, the estimates differ substantially among different studies (35-41).

Herein, I attempt to formulate a new evolutionary framework called *generalized background selection* (Table 1), which includes two major types: (*i*) *negative background selection*, exemplified by the conversional background selection at closely-linked sites (5); and (*ii*) *positive background selection*, exemplified by the selective sweep at closely-linked sites (3). A new diffusion-limit model with two killing function (42) is then developed: the killing function for negative background selection measures the rate for the stochastic trajectory of a fixation process of a mutation to be randomly stopped, whereas the killing function for positive background selection measures the rate for that of a loss process of a mutation to be randomly stopped. The relationship between intra-population diversity and interspecies divergence is then derived, which can be applied to analyze the patterns of inter-species divergence and intra-species diversity in low-recombination regions, while the recombination rate is treated as a biological variable underlying the strength of killing functions. The current study focuses on two fundamental issues: first, which one, selective sweeps or (negative) background selection, is more dominant (7, 24, 27-29, 43); and second, whether either one of them is sufficient to explain the observed diversity and divergence pattern (35-41).

**Table 1.**
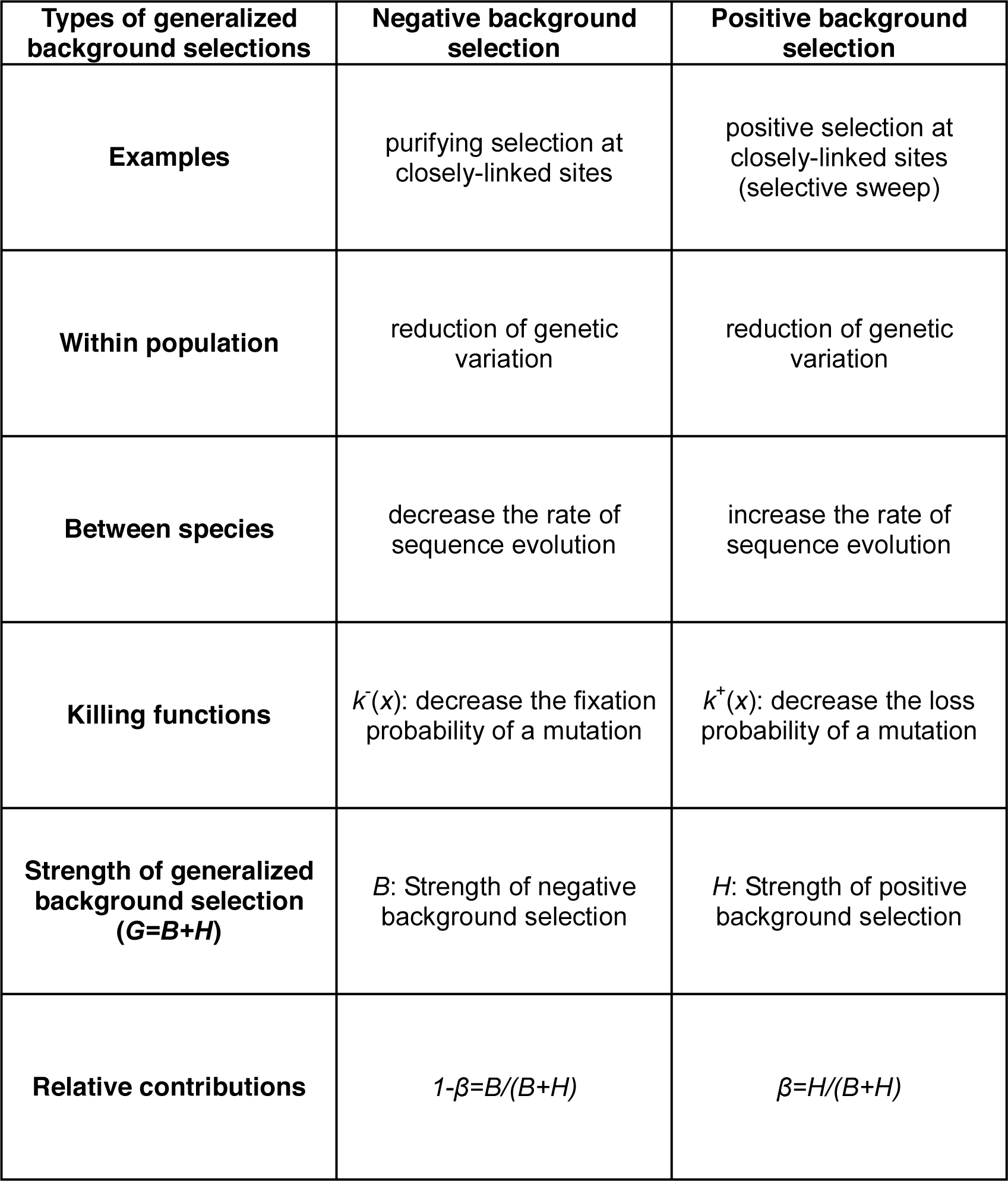
A summary for the conceptual framework of the generalized background selection

## Results

### Diffusion model under the generalized background selection

A conceptual framework of generalized background selection is introduced as follows (Table 1). (*i*) *Negative background selection:* for instance, purifying selection against deleterious mutations may reduce intra-species diversity and interspecies divergence at closely-linked sites (5, 44-47). Since deleterious mutations are prevalent in the genome, many authors argued that (negative) background selection should be part of the basic model of genome variation and evolution (26, 27, 48, 49). (*ii*) *Positive background selection*: for instance, neutral or nearly neutral mutations can be rapidly fixed by a few favorable mutations in the surrounding chromosome region (selective sweeps), resulting in a considerable reduction of genetic variation (3) and an elevated rate of molecular evolution at closely-linked sites. Stephan (11) reviewed different inference methods that have been developed to detect selective sweeps and to localize the targets of directional selection in the genome.

The diffusion-limit model with two killing functions (42) is utilized to model the effects of generalized background selection. Let *x* be the initial frequency of a mutation in the population. (*i*)The killing function associated with negative background selection, denoted by *k*^*-*^(*x*), is the rate for the stochastic trajectory of a fixation process to be randomly stopped, decreasing the fixation probability of a mutation. (*ii*) The killing function associated with positive background selection, denoted by *k*^+^(*x*), is the rate for the stochastic trajectory of a loss process to be randomly stopped, decreasing the loss probability of a mutation from the population (or, indirectly, increasing the fixation probability). In the current study, both killing functions are specified in the form of *k*^-^ (*x*)=*bx*(1-*x*) and *k*^+^(*x*)=*hx*(1-*x*), where *b* and *h* are the coefficients of negative and positive background selections, respectively. These forms of killing functions are not only mathematically convenient, but also reflect the fact that the generalized background selection is effective for mutations with low frequencies.

Let *u*(*x*) be the fixation probability of a mutation with the initial frequency *x*. As shown in Materials and Methods, the Kolmogrov backward equation of *u*(*x*) with two killing functions specified above is given by

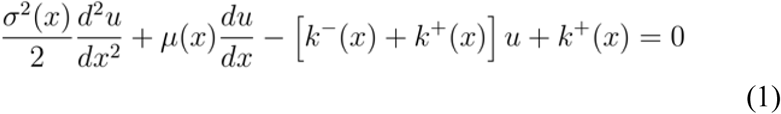

with boundary conditions *u*(0)=0 and *u*(1)=1. The mean μ(*x*) and variance σ^2^(*x*) parameters can be determined under the Wright-Fisher model (see below).

Next we consider intra-species genetic diversity. In the case of no over-dominance, any mutation that appears in a finite population is either ultimately lost or fixed. The effects of generalized background selection make the maintenance of intra-species diversity difficult, because both killing functions tend to reduce the genetic diversity by increasing the chance of a mutation to be either fixed or lost. Nevertheless, under the steady flux of new mutations over many generations, a balance will be reached between production of new mutants and their random loss or fixation. Under this statistical equilibrium there is a stable frequency distribution at different sites in which mutations are neither fixed nor lost (50). Denote the stable frequency of mutations by *p*. Given the initial frequency *x*, let *J(x)=E*[*2p(1-p)*] be the expected heterozygosity of a nucleotide site. As shown in Materials and methods, the steady-flux model with two killing functions claims that *J*(*x*) satisfies the following backward equation

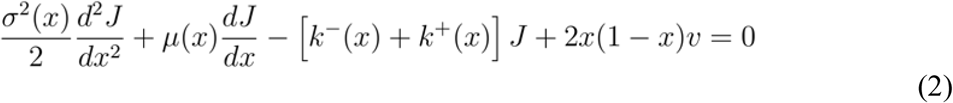

where *v* is the mutation rate; the boundary conditions of Eq.(2) are *J*(0)=*J*(1)=0.

### Neutral interspecies divergence under generalized background selection

An intriguing question is to what extent the prediction of the neutral theory, i.e., the rate of neutral evolution (λ) equals to the mutation rate (*v*), could be affected by the effects of (negative) background selections and selective sweeps (41, 51-53). To this end, we obtained the fixation probability *u*(*x*) given by solving Eq.(1) under the neutral model (see Materials and Methods). Note that the rate (λ) of molecular evolution is defined by the amount of new mutations 2*N*_*e*_*v* multiplied by the fixation probability *u*(*x*) with the initial frequency *x*=1/2*N*_*e*_; where *N*_*e*_ is the effective population size. Let *H*=4*N*_*e*_*h* be the intensity of positive background selection, *B*=4*N*_*e*_*b* be the intensity of negative background selection, and *G=B*+*H* be the intensity of generalized background selection, respectively (Table 1). Putting together, one can show

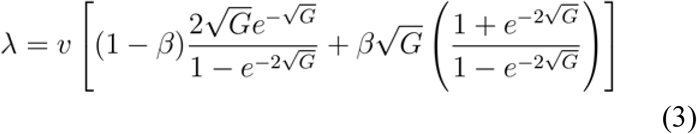

where *β*=*H/(B*+*H)=h/(b*+*h)* is the proportion of positive background selection (selective sweeps), or 1-*β* is that of negative background selections. Eq.(3) shows that, in addition to the mutation rate (*v*), the rate of neutral rate (λ) can be affected by the strength of generalized background selection (*G*) and the proportion of positive background selection (*β*). While the first part on the right hand of Eq.(3) is the rate component associated with negative background selection, the second one is that associated with positive background selection. It appears that Eq.(3) is reduced to λ=*v* (54) when *G*=0.

**Fig.1** panel A presents the plotting of the (neutral) rate-mutaion ratio λ/*v* against the strength (*G*) of generalized background selection under various proportion (*β*) of positive background selection. (*i*) When *β*=0 (i.e., no positive background selection, and so *G*=*B*), the rate of neutral evolution (λ) is always less than the mutation rate (*v*); for instance, λ is about 85%, 55% and 27% of the mutation rate (*v*) when *B*=1, 4 and 10, respectively. Indeed, a very strong negative background selection would virtually cease the neutral divergence between species, i.e., λ≈*0*. (*ii*) By contrast, when *β*=1 (i.e., complete positive background selection, and so *G=H*), the rate (λ) of neutral evolution is always larger than the mutation rate (*v*); for instance, λ is about 131%, 204% and 316% of the mutation rate (*v*) for *H*=1, 4 and 10, respectively. When *H*>1, the rate of neutral evolution is asymptotically linear with the squared root of *H*. (*iii*) In the general case of 0<*β*<1, a numerical analysis of Eq.(3) indicates a critical value *β*_*c*_≈0.334. When *β*<*β*_*c*_, a weak positive background selection, the neutral rate decreases from λ=*v* at *G*=0 with the increasing of *G* until approaching to a minimum; since then λ increases with the increasing of *G*, ultimately toward λ>*v*. On the other hand, when *β*>*β*_*c*_, an intermediate or strong positive background selection, the neutral rate λ increases with the increase of *G* such that λ>*v* always holds.

**Fig.1.**
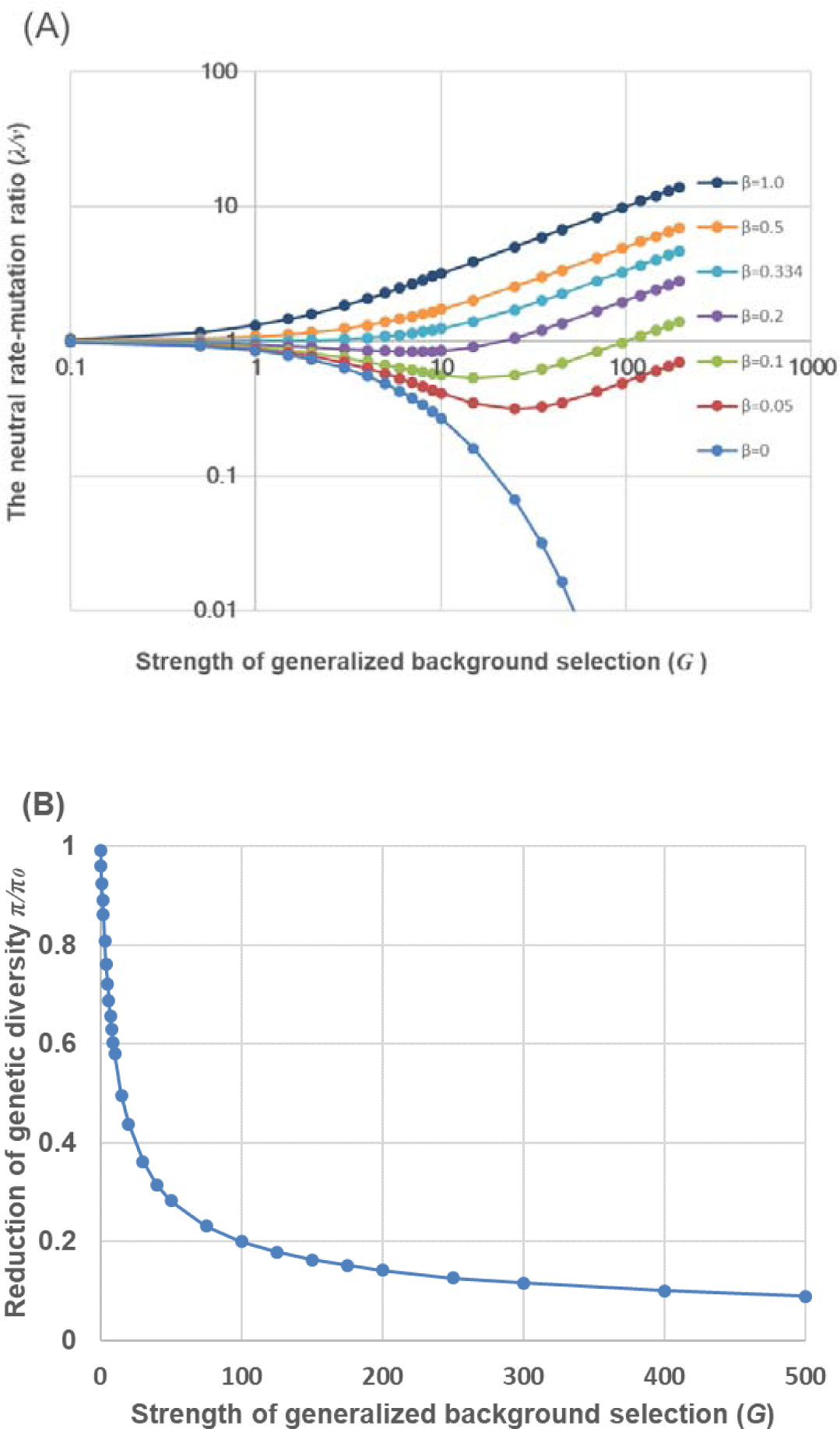
(A) The ratio of neutral evolutionary rate to mutation rate (λ/*v*) plotted against the strength (*G*) of generalized background selection, given the proportion of positive background selection *β*=0, 0.05, 0.1, 0.2, 0.5 or 1.0, respectively. Note that there exists a critical value *β*_*c*_≈0.334. When *β*<*β*_*c*_, the ratio λ/*v* decreases toward the area of λ/*v*<1 with the increasing of *G* until approaching to a minimum; since then λ/*v* increases with the increasing of *G*, ultimately toward the area of λ/*v*>1. When *β*>*β*_*c*_, the ratio λ/*v* increases with the increase of *G* such that λ/*v>*1 holds always. (B) The ratio of neutral genetic diversity (*π/π*_*0*_) plotted against the strength (*G*) of generalized background selection. Here *π* is the expected neutral intra-population diversity under the generalized background selection, and *π*_*0*_ is that with no generalized background selection.

### Neutral genetic variation under generalized background selection

In contrast to the interspecies divergence, both negative and positive selections contribute to the reduction of intra-species genetic diversity (Table 1), which can be derived as follows. One can solve Eq.(2) to obtain the expected neutral heterozygosity of a site, *J*(*x*), in the case of neutrality, i.e., μ(*x*)=0 and σ^2^=*x*(1-*x*)/2*N*_*e*_ (Materials and Methods). As the expected neutral genetic diversity (*π*) of a site is given by *π*=2*N*_*e*_*J*(*x*) at *x*=1/2*N*_*e*_, we show that the expected neutral genetic diversity under the generalized background selection is given by

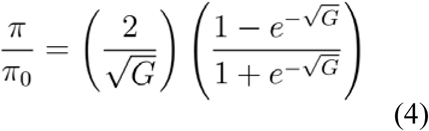

where *π*_*0*_=4*N*_*e*_*v* is the neutral genetic diversity without generalized background selection. It appears that the ratio of neutral genetic diversity (*π/π*_*0*_) depends on a single parameter *G*, the strength of generalized background selection; *π/π*_*0*_<1 always holds, and *π/π*_*0*_=1 only when *G*=0 (**Fig.1 panel B**). Since *G=4N*_*e*_*g*, where *g=b*+*h* is the coefficient of generalized background selection, Eq.(4) indicates that reduction of intra-population neutral diversity by the generalized background selection becomes severe in a large population (in a scale of the squared root of *N*_*e*_), whereas this reduction can be compromised by genetic drifts in a small population.

### Statistical analysis of chromosome regions with low recombination

A number of studies (26, 27, 33, 34, 55) indicated a dominant role of (negative) background selection on the reduction of genetic diversity in regions with low recombination. However, it remains highly controversial whether it is sufficient to explain the observed intra-population diversity and interspecies divergence, without invoking selective sweeps. It appears that Eq.(3) and Eq.(4) together provide a straightforward procedure to address this key issue. **Fig.2** shows the λ/*v*-*π/π*_*0*_ curve for different values of *β*, the proportion of positive background selection. Impressively, for genes with considerable reduced genetic diversity, say, *π/π*_*0*_<0.5, the degree of interspecies divergence (λ/*v*) is highly sensitive to the value of *β*.

**Fig.2.**
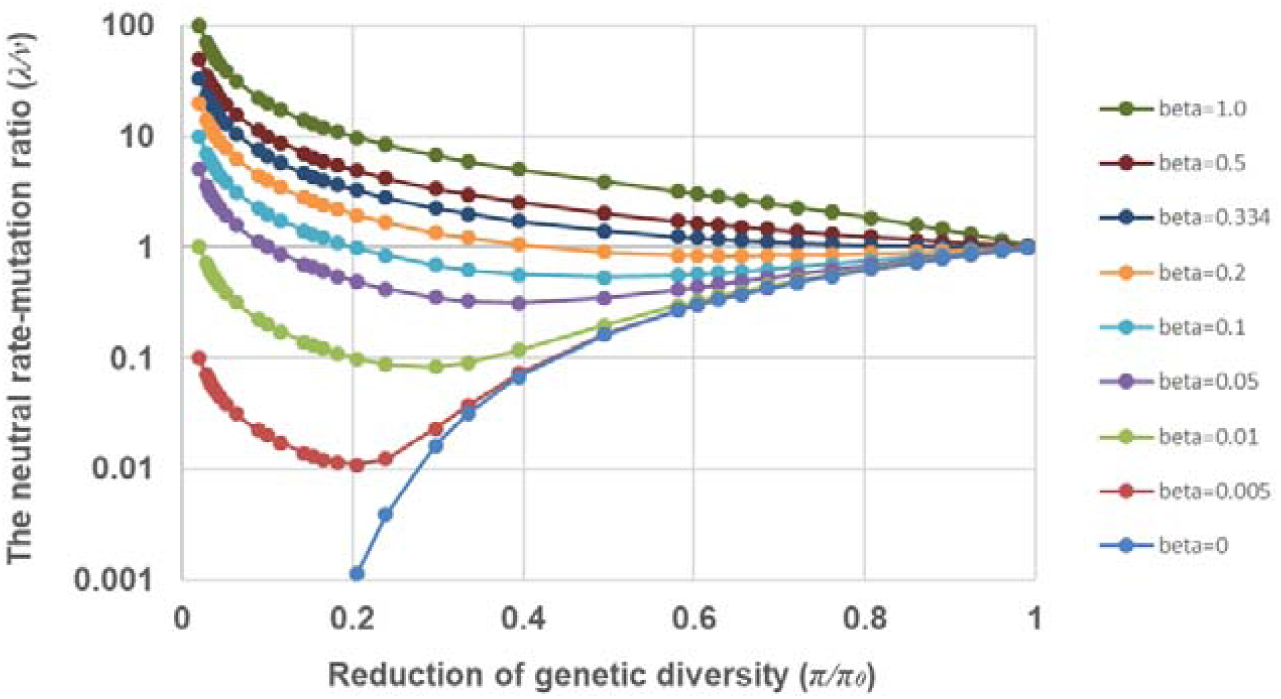
The λ/*v*-*π/π*_*0*_ curve, while the proportion of positive background selection *β*=0, 0.005, 0.01, 0.05, 0.1, 0.2, 0.5 and 1.0, respectively.

#### Statistical procedure

By contracting chromosome regions with low recombination to those with free recombination, one can design a simple computational procedure to estimate *G* and *β*. Suppose that we have two sets of genes: (*i*) genes are located in chromosome regions with free recombination, with the neutral intra-species diversity denoted by *π*_*0*_, and the interspecies neutral divergence denoted by *d*_*S0*_. And (*ii*) genes are located in chromosome regions with low recombination, with neutral intra-species diversity denoted by *π*, and the interspecies neutral divergence denoted by *d*_*S*_. After calculating the *π/π*_*0*_ ratio, the strength of generalized background selection (*G*) for genes in low-recombination regions is obtained by numerically solving Eq.(4). Next, one can calculate the divergence ratio *d*_*S*_/*d*_*S0*_ as a proxy to the λ/*v* ratio in Eq.(3), allowing to estimate *β* after replacing the parameter *G* by its estimate.

#### Non-crossover regions (NC) of D. melanogaster

Campos et al. (27) used next-generation DNA sequence data of a population of *D. melanogaster* to compare the intra-population diversity and interspecies divergence across the whole genome. They analyzed 268 genes located in five independent heterochromatic regions that lack crossover (‘non-crossover regions’, NC) of *D. melanogaster*, contrasting to genes located in the crossover regions (AC short for autosomes and XC for X-chromosome). For each gene group, the mean sequence divergence between *D. melanogaster* and *D. yakuba* was calculated.

We reanalyzed Campos et al. (27)’s data by computing the ratio *π*/*π*_*0*_: *π* is from synonymous diversity in each of five NC regions (and the pooled), and *π*_*0*_ is from that in AC or XC, respectively. According to Eq.(4), it is straightforward to estimate *G*, the strengths of generalized background selection. **Table 2** shows that the estimate of *G* ranges from 0.40×10^2^∼12.2×10^2^. This variation of estimated *G* may reflect different recombination rates among NC regions, but the 95% CI (confident interval) of estimated *G* is broad. Next we estimated *β* (the proportion of positive background selection) by Eq.(3), where the ratio λ*/v* is calculated by the interspecies synonymous distances in NC regions and AC/XC regions, respectively (Table 2). It is impressive that the range of estimated *β* among five NC regions is narrow, ranging from 2.48% to 15.1%.Moreover, we used the pooled NC data to statistically test the null hypothesis of no positive background selection (*β*=0). In the case AC as crossover regions, the estimated *β*=8.06%, with 95% CI from 3.51% to 12.0%, and virtually the same results in the AC case (Table 2). Therefore, the pooled NC region analysis suggests that, to explain the pattern of genetic diversity and divergence, a weak but significant positive background selection (selective sweeps) in NC regions is required.

**Table 2.**
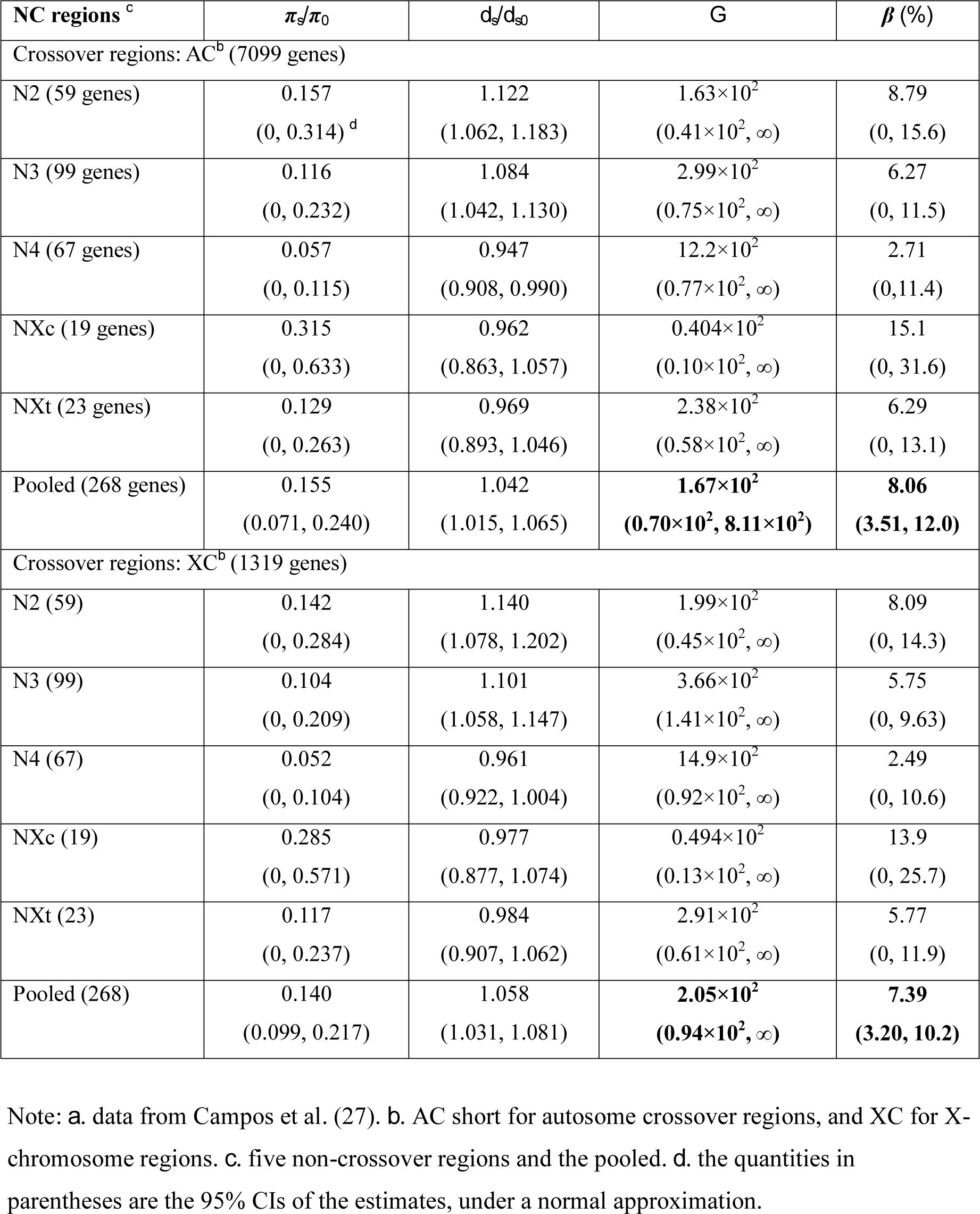
Statistical analysis of *Drosophila* population genomics data^*a*^.

| NC regions<sup>c</sup> | $\pi_s/\pi_0$ | $d_s/d_{s0}$ | $G$ | $\beta$ (%) |
| --- | --- | --- | --- | --- |
| Crossover regions: AC<sup>b</sup> (7099 genes) |  |  |  |  |
| N2 (59 genes) | 0.157<br>(0, 0.314)<sup>d</sup> | 1.122<br>(1.062, 1.183) | $1.63 \times 10^2$<br>($0.41 \times 10^2$, ∞) | 8.79<br>(0, 15.6) |
| N3 (99 genes) | 0.116<br>(0, 0.232) | 1.084<br>(1.042, 1.130) | $2.99 \times 10^2$<br>($0.75 \times 10^2$, ∞) | 6.27<br>(0, 11.5) |
| N4 (67 genes) | 0.057<br>(0, 0.115) | 0.947<br>(0.908, 0.990) | $12.2 \times 10^2$<br>($0.77 \times 10^2$, ∞) | 2.71<br>(0, 11.4) |
| NXc (19 genes) | 0.315<br>(0, 0.633) | 0.962<br>(0.863, 1.057) | $0.404 \times 10^2$<br>($0.10 \times 10^2$, ∞) | 15.1<br>(0, 31.6) |
| NXt (23 genes) | 0.129<br>(0, 0.263) | 0.969<br>(0.893, 1.046) | $2.38 \times 10^2$<br>($0.58 \times 10^2$, ∞) | 6.29<br>(0, 13.1) |
| Pooled (268 genes) | 0.155<br>(0.071, 0.240) | 1.042<br>(1.015, 1.065) | **$1.67 \times 10^2$**<br>**($0.70 \times 10^2$, $8.11 \times 10^2$)** | **8.06**<br>**(3.51, 12.0)** |
| Crossover regions: XC<sup>b</sup> (1319 genes) |  |  |  |  |
| N2 (59) | 0.142<br>(0, 0.284) | 1.140<br>(1.078, 1.202) | $1.99 \times 10^2$<br>($0.45 \times 10^2$, ∞) | 8.09<br>(0, 14.3) |
| N3 (99) | 0.104<br>(0, 0.209) | 1.101<br>(1.058, 1.147) | $3.66 \times 10^2$<br>($1.41 \times 10^2$, ∞) | 5.75<br>(0, 9.63) |
| N4 (67) | 0.052<br>(0, 0.104) | 0.961<br>(0.922, 1.004) | $14.9 \times 10^2$<br>($0.92 \times 10^2$, ∞) | 2.49<br>(0, 10.6) |
| NXc (19) | 0.285<br>(0, 0.571) | 0.977<br>(0.877, 1.074) | $0.494 \times 10^2$<br>($0.13 \times 10^2$, ∞) | 13.9<br>(0, 25.7) |
| NXt (23) | 0.117<br>(0, 0.237) | 0.984<br>(0.907, 1.062) | $2.91 \times 10^2$<br>($0.61 \times 10^2$, ∞) | 5.77<br>(0, 11.9) |
| Pooled (268) | 0.140<br>(0.099, 0.217) | 1.058<br>(1.031, 1.081) | **$2.05 \times 10^2$**<br>**($0.94 \times 10^2$, ∞)** | **7.39**<br>**(3.20, 10.2)** |

Note: *a.* data from Campos et al. (27). *b.* AC short for autosome crossover regions, and XC for X-chromosome regions. *c.* five non-crossover regions and the pooled. *d.* the quantities in parentheses are the 95% CIs of the estimates, under a normal approximation.

#### Pericentromeric regions in Soybean (Glycine max)

Du et al. (43) calculated synonymous distances (*K*_*S*_) of genes between soybean (*Glycine max*) and its annual wild relative, *Glycine soja*. They compared the mean *K*_*S*_ (*K*_*S,arm*_) located in a chromosomal arm (high recombination rate) and those (*K*_*S,peri*_) in a pericentromeric region (low recombination rate) in three genomic datasets: (*i*) high-confidence genes annotated in the soybean reference genome; (*ii*) singletons (single-copy genes) from high-confidence genes; and (*iii*) WGD (whole genome duplication) duplicate pairs, each of which has one copy in a chromosomal arm and the other one in a pericentromeric region. The ratio *K*_*S,peri*_*/K*_*S,arm*_ is 0.820, 0.764, and 0.731 for the three datasets, respectively. It has been roughly estimated that the ratio of synonymous diversity in pericentromeric regions to chromosomal arms within soybean population, denoted by*π*_*S,peri*_*/π*_*S,arm*_, is about 0.19 (56). In this case, one may obtain the estimate of strength of generalized background selection (*G*) as 110.3, and the estimates of the proportion of positive background selection (*β*) as 7.73%, 7.14%, and 6.81% for three datasets, respectively.

### Empirical relationship of *G* with the rate of recombination (*r*)

Numerous analyses have established a well-known positive correlation between the genetic diversity (*π*) and the recombination rate (*r*) (6-11, 14). When the current model is applied to chromosome regions with different *r* values, we expect an inverse relationship between the strength of generalized background selection (*G*) and the recombination rate, as demonstrated by Eq.(4) that genetic diversity (*π*) is inversely determined by *G* (Fig.1 panel B).

We analyzed the inter-species divergence and intra-species variation in different human chromosome regions with different recombination rate measured by cM/Mb; data from Nachman (24). **Fig.3 panel A** shows the mean strength of generalized background selection (*G*) in genes located in low (cM/Mb<1), middle (1<cM/Mb<2) and high (cM/Mb>2) recombination regions, respectively. A strong generalized background selection in low recombination regions has been observed. Meanwhile, **Fig.3 panel B** shows the mean proportion of positive background selection (*β*) in genes located in low, middle and high recombination regions, respectively. Interestingly, the estimate of *β=*8.9% in human low recombination region is very similar to that in fruitfly (∼8%) or soybean (7%-8%).

**Fig.3.**
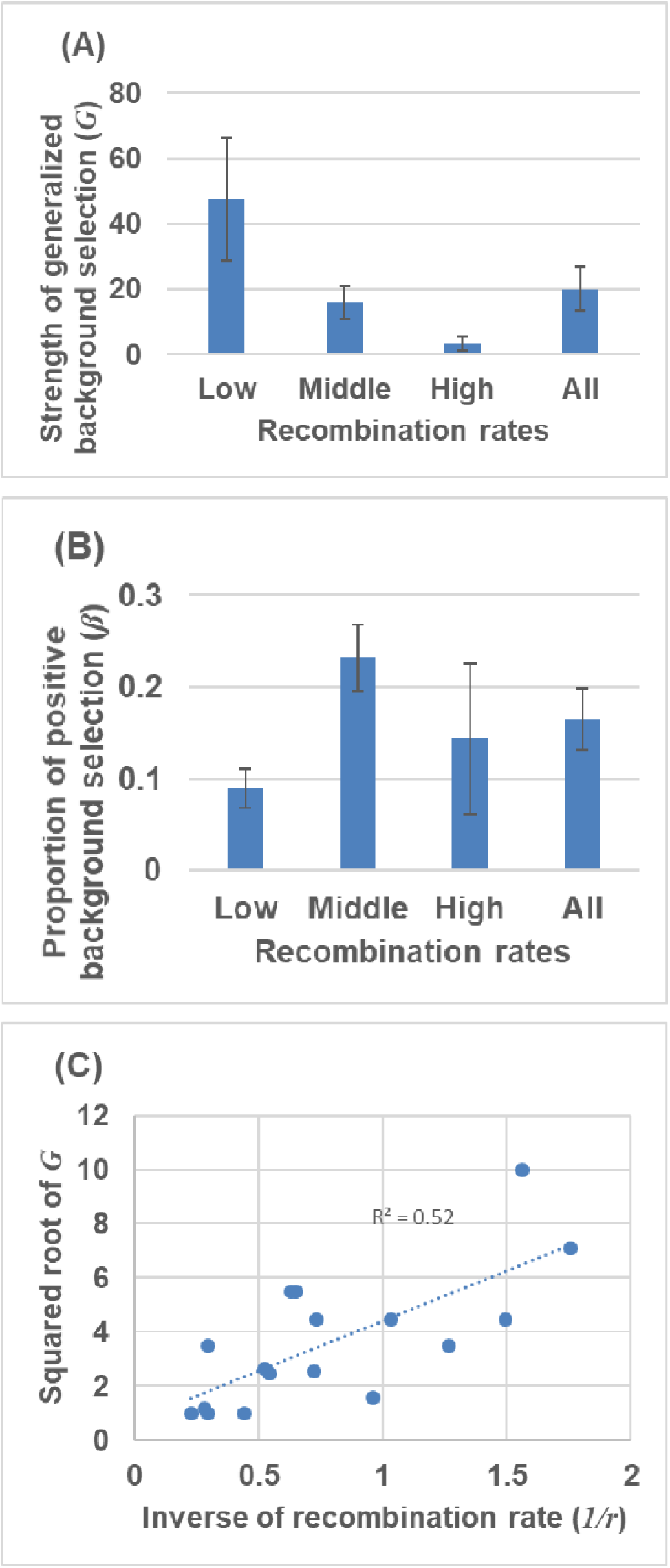
Analysis of divergence and variation in human chromosome regions with different recombination rate (*r*, measured by cM/Mb); data from Nachman (24). (A) The mean strength of generalized background selection (*G*) in genes located in low (cM/Mb<1), middle (1<cM/Mb<2) and high (cM/Mb>2) recombination regions, respectively, as well as that of all genes. Standard errors are presented. (B) The mean proportion of positive background selection in genes located in low, middle and high recombination regions, respectively, as well as that of all genes. Standard errors are presented. (C) The squared root of *G* estimates plotted against the inverse of *r*, the recombination rate. The coefficient of correlation is 0.73, *P*-value<0.01.

Kim and Stephan (18) showed that for the chromosome region with low recombination rate, the joint effects of deleterious and beneficial mutations on neutral variation can be approximated by

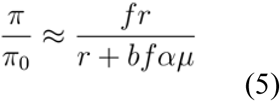

where *r* is the local recombination rate, α is the (positive) selection intensity, μ is the rate of adaptive substitution, *f* describes the reduction of *N*_*e*_ owing to deleterious mutations, and *b* is an empirically-determined constant. While the theoretical derivation of the *G-r* inverse relationship remains challenging, one can establish an empirical one by equating Eq.(5) with Eq.(4); under the assumption of *G>*1, it is approximated by

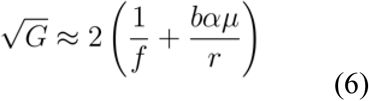

Indeed, **Fig.3 panel C** shows a rough linear relationship between the squared-root of *G* and the inverse of *r*; the coefficient of correlation is 0.73, *P*-value<0.01.

## Discussion

Based on the diffusion-limit model with two killing functions, I have formulated a framework of generalized background selection that has two predictions: (*i*) reduction of intra-species neutral diversity is inversely determined by a single parameter (*G*), the strength of generalized background selection that combines both negative or positive effects; and (*ii*) the rate of neutral divergence between species decreases with *G*, but increases with the proportion (*β*) of positive background selection. Many studies attempted to determine the relative impact of selective sweeps to (negative) background selections on closely-linked sites. For instance, Pouyet et al. (33) concluded that the negative background selection influences as much as 85% of the genetic variants of the human genome, and Campos et al. (27) claimed that a strong selective sweep was unlikely in non-crossover regions of *Drosophila*. Our case-studies in three organisms (fruitfly, soybean and human) showed that the proportion (*β*) of positive background selection (selective sweeps) in chromosome regions with low recombination rate is statistically significant, though on average, less than 10%.

Compared to previous work (6-11), the new model may have some advantages. First, it provides a straightforward approach to data analysis without oversimplification about the selection themes, as shown by Eqs.(3) and (4). The role of recombination rate (*r*) is elaborated by its inverse relationship with the strength of generalized background selection (*G*). Second, the new model provides a biologically intuitive explanation why the mechanism of selection sweeps is essential. Suppose *G*=167 as estimated from the (pooled) NC regions of *Drosophila*. If the (negative) background selection is the only mechanism underlying the reduced genetic diversity, i.e., *β*=0, the rate of neutral divergence between species calculated by Eq.(3) would be as low as 10^−5^ of that in crossing-over regions, that is, the Muller’s ratchet (57) virtually ceases the pace of evolution. Surprisingly, if we assume *β*=0.02, i.e., a very small portion of selective sweeps, the rate of neutral divergence is only as low as 26% of that with free recombination (Fig.1 panel A). Third, one may design a new Bayesian approach for the detection of sites targeted by positive selection: for a given chromosome region with dramatically reduced diversity, a high level of sequence divergence between closely-related species may indicate the role of selective sweeps. Finally, our analysis may shed some lights on the recent debates on Neutral Theory (12, 13). If the proportion (*β*) of selective sweeps estimated from chromosome regions with low recombination is generally applicable to the whole genome, one may conclude that neutral or nearly neutral selection dominates the genome-wide variation and evolution, while approximately 8% of mutations may be subject to positive selection.

An immediate extension of the current analysis is to include non-neutral sites such as nonsynonymous sites (27, 33, 43, 51). It has been shown that the interplay between *G, β* and *S* (the strength of ‘direct’ selection on the site under study) reveals more sophisticated evolutionary scenarios (Gu, unpublished results). There are several challenges remaining. First, the detailed structure of the inverse relationship between *G* and the recombination rate is desirable for both theoretical and empirical studies. Second, as time-dependent *N*_*e*_ changes may affect the estimation of selection sweeps (7, 9, 11, 20, 41), the effect of constant *N*_*e*_ assumed in the current model needs to be investigated. Third, it has been shown that the MacDonald-Kreitman (MK) test could be affected by both background selection and selective sweeps (61). An interesting question is to what extent the key parameters (*G* and *β*) may influence the MK test. And forth, several factors, such as GC content and biased gene conversion, can also influence the reduced intra-population genetic reduction (25, 58, 59). It is important to remove those factor before the current method is applied. We will address these challenges in the future study, with the help of extensive computer simulation studies such as SLim (60).

## Materials and Methods

### Datasets

We used the genome-wide genetic diversity profiles of *D. melanogaster* provided by Campos et al. (27). They compiled 268 genes, which located in five independent heterochromatic regions that lack crossover (‘non-crossover regions’, NC) of *D. melanogaster*, contrasting to the crossover regions (AC short for autosomes and XC for X-chromosome). For each chromosome region under study, the sequence divergence with *D. yakuba* is also calculated.

Soybean (*Glycine max*) was domesticated from its annual wild relative, *Glycine soja*, about 5000 years ago. One striking feature of the soybean genome is that ∼57% of the genomic sequence occurs in recombination-suppressed heterochromatic regions surrounding centromeres (referred to as pericentromeric regions). We used three genomic datasets compiled by Du et al. (43): (*i*) high-confidence genes (27571) annotated in the soybean reference genome; (*ii*) 12,994 singletons (single-copy genes) from high-confidence genes; and (*iii*) 2439 WGD (whole genome duplication) duplicate pairs, each of which is composed of one copy in a chromosomal arm and the other one in a pericentromeric region.

Human dataset including 17 genes are obtained from Nachman (24) for which the sample size is greater than ten. For each gene, the genetic diversity (*π*) in the human population, the sequence divergence with chimpanzees, and the recombination rate (in the human genome, measured by cM/Mb) are available.

### Kolmogrov backward equations with two killing functions

#### Fixation probability

Let *u*(*x*) be the probability of an allele to be ultimately fixed in the population, given the initial allele frequency (*x*). By the standard diffusion theory, *u*(*x*) satisfies the following Kolmogrov backward equation

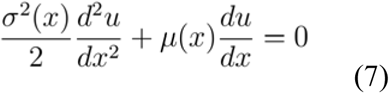

with the boundary conditions *u*(0)=0 and *u*(1)=1. Second we consider the backward equation of *u*(*x*) with a killing function associated with the negative background selection, *k*^*-*^(*x*), which is the rate for the stochastic trajectory of fixation process to be randomly stopped. It appears that *k*^*-*^(*x*) decreases the fixation probability of a neutral or nearly-neutral mutation. Karlin and Taylor (42) showed that in this case *u*(*x*) follows

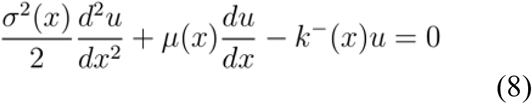

Next we consider the backward equation of *u*(*x*) with a killing function *k*^+^(*x*) associated with the positive background selection. Let *u*\*(*x*)=1-*u*(*x*) be the ultimate loss probability of an allele. The killing function *k*^+^(*x*) is then defined by the rate for the stochastic trajectory of loss process to be randomly stopped. Hence, *k*^+^(*x*) tends to decrease *u*\*(*x*) and so to increase the fixation probability *u*(*x*)=1-*u*\*(*x*). One can show that *u*\*(*x*) satisfies the backward equation similar to Eq.(8) except for *k*^+^(*x*), with the boundary conditions *u\**(0)=1 and *u\**(1)=0. After replacing *u*\*(*x*) by *u*(*x*)=1-*u*\*(*x*), we obtain

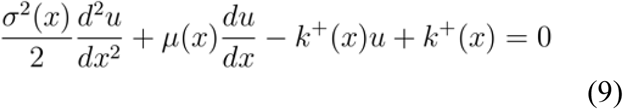

Finally, we derive the backward equation under the joint effects of two killing functions under the assumption that the negative and positive background selections are two independent mechanisms. It follows that the Kolmogrov backward equation for *u*(*x*) can be formulated by combining Eqs.(8) and Eq.(9), resulting in Eq.(1).

#### Intra-population genetic diversity

Let *J(x)=E*[*2p(1-p)*] be the expected heterozygosity of a nucleotide site. Under the standard steady-flux model, it is known that *J*(*x*) satisfies the following backward equation

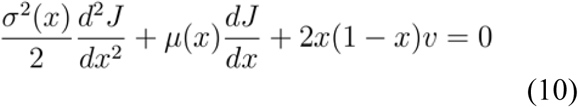

Since both killing functions tend to reduce the genetic diversity by increasing the chance of a mutation to be either fixed or lost, the sum of two killing functions, *k*^+^(*x*)+*k*^*-*^(*x*), can be considered as a single combined killing function. According to Karlin and Taylor (42), we obtain Eq.(2).

#### Derivation of Eq.(3) and Eq.(4)

Under the selectively neutral model with two killing functions, we have μ(*x*)=*0*, σ^2^=*x*(1-*x*)/2*N*_*e*_, *k*^*-*^(*x*)=*bx*(1-*x*), and *k*^+^(*x*)=*hx*(1-*x*). Eq.(1) can be simplified as follows

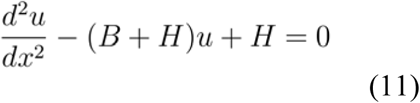

where *B=4N*_*e*_*b, H=4N*_*e*_*h* and *G=B*+*H*. The general solution of Eq.(11) can be written as 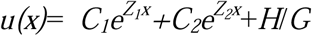, where *Z*_*1*_*=G*^*1/2*^ and *Z*_*2*_*=-G*^*1/2*^ and constants *C*_*1*_ and *C*_*2*_ are determined by the boundary conditions *u*(0)=0 and *u*(1)=1. When *x* is small, it is convenient to use the approximations exp(*Z*_*1*_*x*)≈1+*Z*_*1*_*x* and exp(*Z*_*2*_*x*)≈1+*Z*_*2*_*x*, respectively. After *u*(*x*) is obtained, it is straightforward to have Eq.(3), with a new parameter *β*=*H*/*G*. In the same manner, under the neutral model, Eq.(4) can be simplified as follows

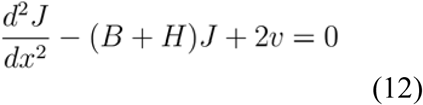

which can be easily solved under the conditions of *J*(0)=*J*(1)=0.

### Statistical evaluation of parameter estimation

The 95% confidence intervals (CI) for estimated *G* and *β* from *Drosophila* genome dataset can be approximately determined as follows. Based on the 95% CIs for synonymous diversities and distances provided by the original authors (27), we simulated a joint sampling density of these measures under the normal assumption, which can be used to empirically determine the 95% CIs of the estimates *G* and *β*.

## Acknowledges

The author is grateful to all members of my research group for constructive comments in the early version of this manuscript.

